# The role of noise in the tumour-immune interactions reveals memoryless shifting response

**DOI:** 10.1101/2024.05.12.593781

**Authors:** Yamen Alharbi

## Abstract

In the present study, the effects of randomness on the model of the immune and cancer cells is discussed. First, we uncover the existence of a bistable response in this interaction where portion of population are cancerous cells. We then discuss how the individual variation within these cells can affect this bistable response. The dominance duration of the tumour and immune cells are highlighted. We calculated the mean of the elimination time of the tumour cells and how can be affected by some key parameters. The analysis reveals that the switching from a stable state to another has a memoryless property.

## Introduction

Cancer is a complex disease that affects millions of people. It can directly affect individual or indirectly affects economical system. Cancer is occurred when cells growth without control in the body [1]. Cancer can occur in any part such as breast, lung, or prostate. It is one of the leading cause of death worldwide, with significant impacts on individuals, families, and communities [2]. Researchers deal with this condition in order to provide better treatment options, improve survival rates and quality of life for many cancer patients. However, the fight against cancer continues, with ongoing efforts to develop better prevention strategies, early detection methods, and more effective treatments.

Many mathematical models have been introduced to investigate the development of the tumour in the host [3–8]. These models, depending on the level of their complexity, can predict the tumour cells proliferation. They can consider different interacting components, for instance the interactions between the tumour cells, immune cells and healthy cells. Different approaches can be used in modelling these interacting components. Some models are constructed using deterministic approach [2, 9, 30–32] while the other based on the stochastic approach [10–12]. Irina et. al [13] modelled the interaction between healthy and tumour cells using deterministic and stochastic approaches. They show that the system exhibits two stable steady states and are separated by an unstable one. In the stochastic model, they examine the effect of additive random noise (in contrast to what we examine here) on the dynamics of the system using Euler–Maruyama method. They stated that in order to generate a significant change in the dynamical behaviour of the system, the noise intensity has to be relatively high. Another deterministic mathematical model addressing the impact of glucose risk on tumour growth was developed by Alblowy et. al [6]. Three equations each tracked the concentrations of immune, tumour, and normal cells in their model. They demonstrated how consuming more glucose has a negative impact on the concentration of immune cells, which raises the concentration of tumours.

In a recent study, Jawad et. al [3] constructed a deterministic model describing the interaction between tumour and immune cells. The model consists of two different Ordinary Differential Equations (O.D.E.) each of which describe the concentrations of one type of these cells; the model (1) assumes the tumour is treated with chemotherapy. They also examined the role of vitamins to post or enhance the immune cells. They were able to calculate the minimum treatment level in order to eliminate the tumour. In this work, however, they did not take into account the individual variation between the cells in the host. In our work, however, we will investigate the impact of stochastic variation on model (1). We will also investigate the effects of increasing the vitamin intake in generating a bistable response. Since the response of the cells is controlled by chemical reactions, we will use the Stochastic Simulation Algorithm (SSA), which is also known as the Gillespie Algorithm [14, 15], in our investigation. SSA consists of Monte Carlo simulations steps which determines the type of the reaction and which reaction will occur at a given time interval [16].

## Materials and methods

### 1 The model

The deterministic model comprises of two ODEs to track the concentration of the number of the tumour and immune cells. The former type is denoted by 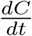 whereas the later is denoted by 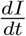 . The production of the tumour cells is assumed to follow a sigmoidal response and the decay rate depends on the presence of the immune cells.

There is a competition behaviour between the tumour cells and immune cells as well. The immune system is enhanced by some vitamin intake *γ* (cf. [3] for details).

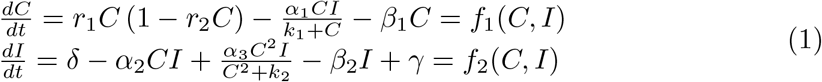

In the original paper [3], the authors studied the deterministic model in details, however, they did not examine the impact of the randomness within the individual cells. Therefore, we will focus our attention on the effect of increasing *δ* on the number of steady states. Using the parameter values from the original paper we can rewrite the steady state equation as a function of *C*. Where *γ* is the controlled parameter.

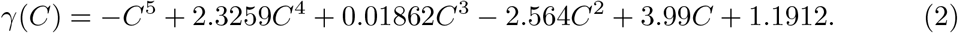

*γ* → ∞ as *C* → −∞ and tends to −∞ as *C* → ∞. Notice that *γ*(0) = 1.1912, therefore there must be at least a positive root for the polynomial. Using Descartes’ rule of sign, equation (2) changes sign three times, hence it can have three positive roots at most. For specific value of *γ* the system can have two steady states plus the trivial one (*C*^∗^ = 0). Hence, the system can have a bistable switch (see Fig (1)). This figure shows that the system has a single positive steady state for low value of *γ* and from this figure it is a stable equilibrium point.

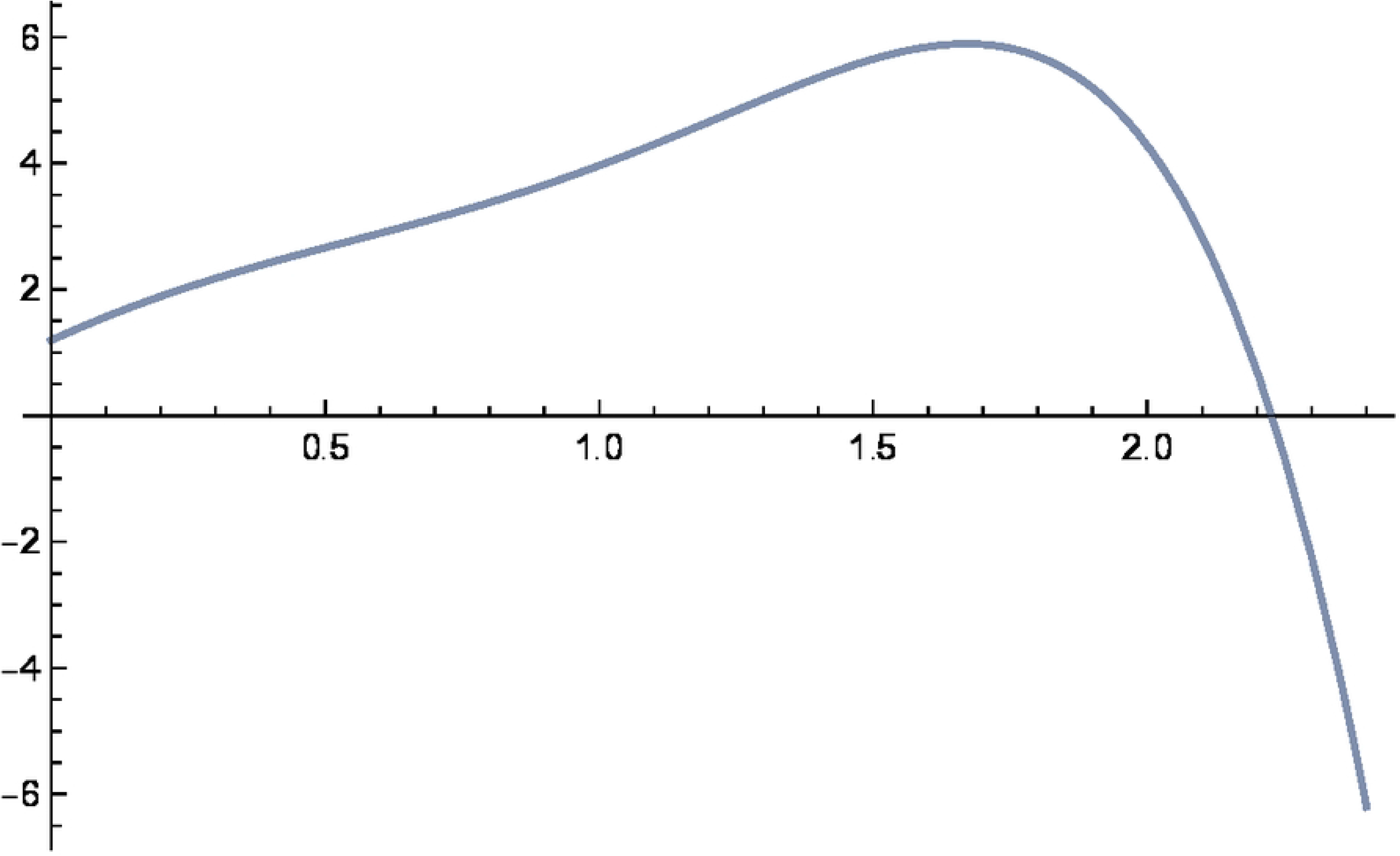

### 2 Bistable switch in the deterministic model

Extensive numerical simulations show that the system has three steady states. Fig (2) has a bifurcation diagram showing the number of steady states as a function of *γ*. In this figure, the red and black curves represent the stable and unstable equilibria, respectively. The system can shift between these two equilibrium points depending on the choices of initial data (see Fig (3)). For low *γ* values, the system trends toward the higher stable steady state (high number of tumour cells) and when the value of *γ* is relatively high, the system trends toward the free tumour state. Between these two thresholds, there is a bistable switch with hysteresis; in this switch, the system remembers its previous state. In other words, the system can jump from higher to lower stable steady state by varying the vitamin intakes, parameter *γ*. Moreover, in this region, the system can have two types of cells, with some cancerous cells coexisting with tumour-free cells. This behaviour was not observed in the original paper. This type of switch arises due the existence of a maximum value in equation (2) (see Fig (1)) [18, 21, 22].

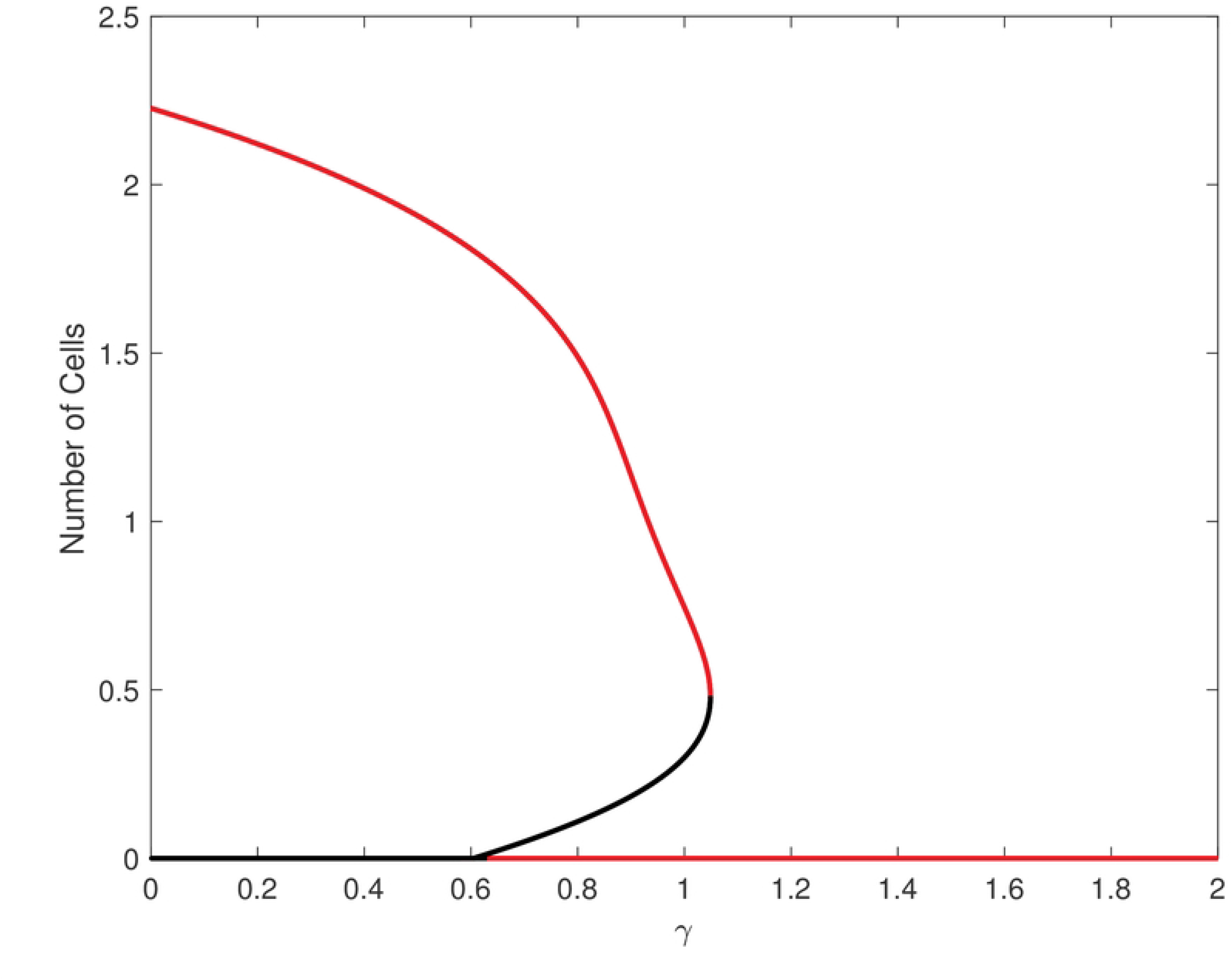

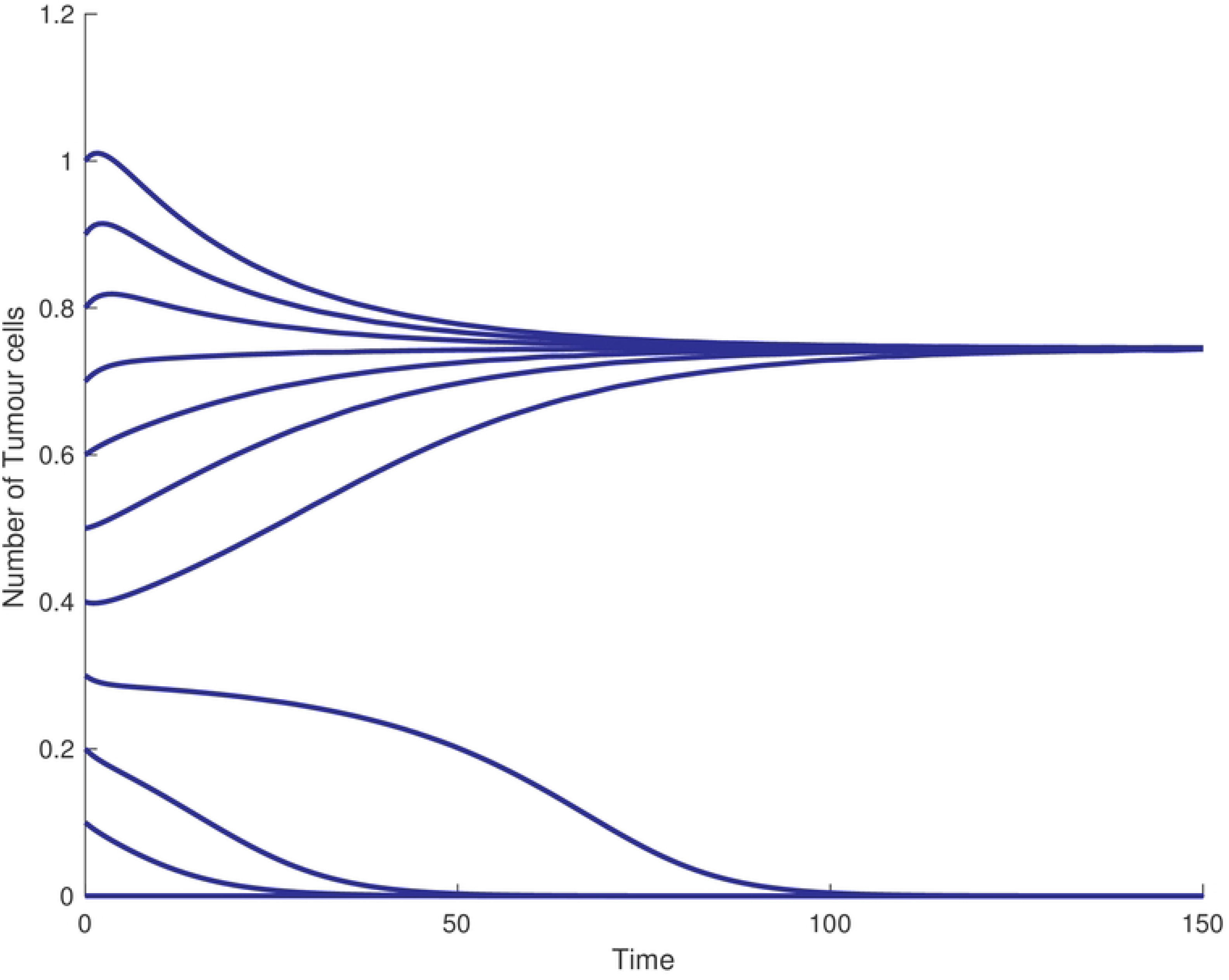

We simulated the system at different initial conditions at the bistable region and observed the following. When the initial data is selected from the region below the unstable steady state, the system tends to the lower stable steady state (tumour-free state). However, when the initial data is selected from the region above the unstable branch, the solution tends to the stable higher basin of attraction (cancer-present steady state).

We can summarise the relationship between chemotherapy treatment with the vitamin intake parameter in a two-parameter diagram (Diagram (4)). This diagram shows that the system can exhibits two regions: bistable and monostable. The diagram also illustrates how the system can generate a bistable switch even when there is no chemotherapy treatment and no vitamin intake. If the system is at the bistable region, increasing *γ* or *β*_1_ will drive the system into the tumour-free state. It is important to note that the system requires a higher dose of chemotherapy (*β*_1_) to eliminate the tumour if there is no vitamin given to the patients than when vitamin is introduced. Interestingly, the dose of the chemotherapy is decreased as we increase the vitamin intake. In other words, with vitamins, the patient requires less chemotherapy than when there is no vitamin intake. This figure summarises the importance of introducing vitamins to the treatment plan. For example, when *γ* = 0, the system requires *β*_1_ ≈ 0.23 in order to push the system into the tumour-free state. However, when *γ* = 0.7, the system requires *β*_1_ ≈ 0.1 in order to reach a tumour-free state. After analysing the behaviour of the system in the deterministic model, we will explore the effects of noise on the system.

#### 2.1 Source of noise in the dynamical system

Stochasticity plays a crucial role in dynamical systems. There are two types of noise or stochastic variations; extrinsic and intrinsic noises. The extrinsic noise is that which arises from an outside source, such as a variation of activation signals. The intrinsic noise refers to the stochastic variations that arise within individual cells [19]. An example of the intrinsic noise is the low copy number of molecules. When the number of molecules is high, the noise cannot have an impact on the system. However, when there is a limited number of molecules, then the noise plays an important role in the fate of the system [16, 20]. In the proceeding sections, we shall discuss the effects of latter type of noise on the behaviour of system (1).

#### 2.2 From deterministic to stochastic system

In the dynamical system, when the number of molecules are high, the randomness does not play role in determining its behaviour. However, when the system has a low number of molecules, stochastic variation can play a crucial role in the response of the system. Throughout the rest of the paper, we will be focusing on the effects of low number of cancer and immune cells in the model. The intrinsic noise is increased as the number of molecules is decreased. Thus, it is essential to include the effects of noise in the models [25]. To simulate the intrinsic noise, we will apply the Stochastic Simulation Algorithm (SSA). This method aims to produce random sampling trajectories which tracks the state of the system as time evolves. Because the reaction occurs randomly, and the number of molecules (cells) in the system is a random variable and its time evolutions depends only on the current state, the Markov process can be applied [26]. The SSA is exact which means the resultant trajectories are exact to the ones generated from the Chemical Master Equation. To assess our investigation, it is essential to convert the model from representing concentration to the number of molecules per cell [25]. Before doing so, we shall construct the reaction diagrams to help us in the conversion from reaction rates to stochastic constant. The diagrams can be written as follows.

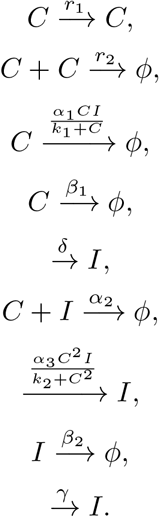

Following Voigt et. al [18] in this regards, we assume that 1 molecule is equal to 1 nM and the rate constants are shown in Table (1).

**Table 1.**
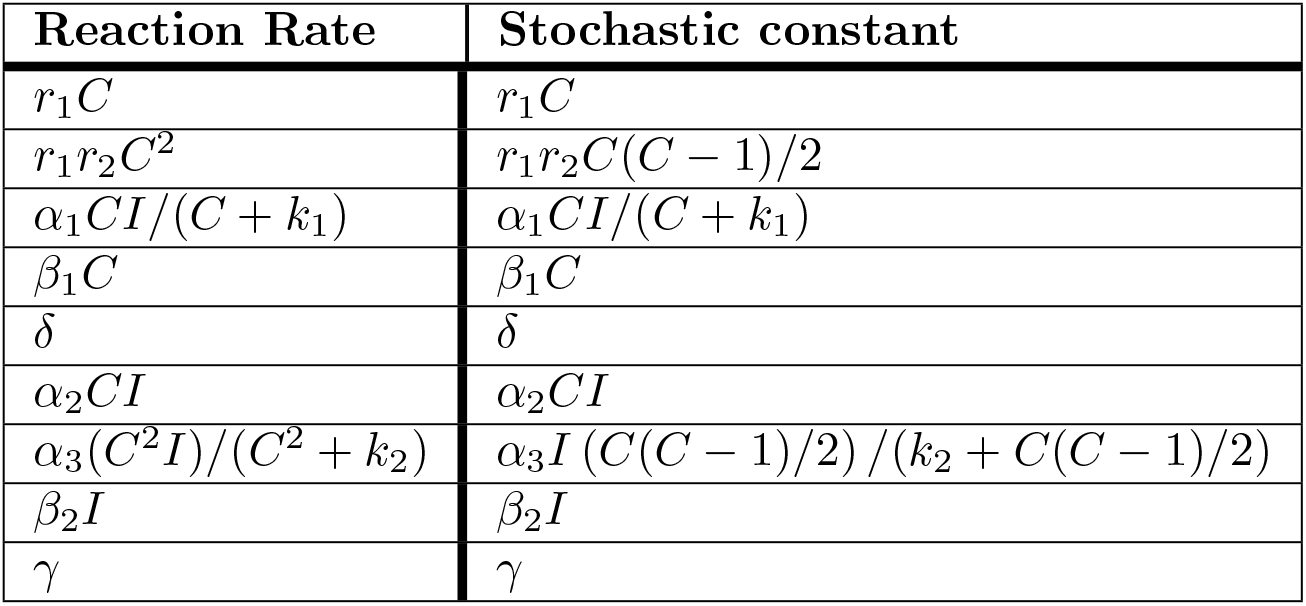
Deterministic rates to stochastic constants conversion.

| Reaction Rate | Stochastic constant |
| --- | --- |
| $r_1 C$ | $r_1 C$ |
| $r_1 r_2 C^2$ | $r_1 r_2 C(C-1)/2$ |
| $\alpha_1 C I / (C + k_1)$ | $\alpha_1 C I / (C + k_1)$ |
| $\beta_1 C$ | $\beta_1 C$ |
| $\delta$ | $\delta$ |
| $\alpha_2 C I$ | $\alpha_2 C I$ |
| $\alpha_3 (C^2 I) / (C^2 + k_2)$ | $\alpha_3 I (C(C-1)/2) / (k_2 + C(C-1)/2)$ |
| $\beta_2 I$ | $\beta_2 I$ |
| $\gamma$ | $\gamma$ |

#### 2.3 Noise destroying the bistable response

To facilities our discussion, we will outline the effect of noise on the monostable and bistable region of Fig 4. We fix the system in the monostable region *γ* = 1.5, and *β*_1_ = 0 and run 10,000 simulations using the SSA. The results are as follows; when the system is at the monostable region, the average number of tumour cells trends to the deterministic stable steady state as illustrated in Fig (5).

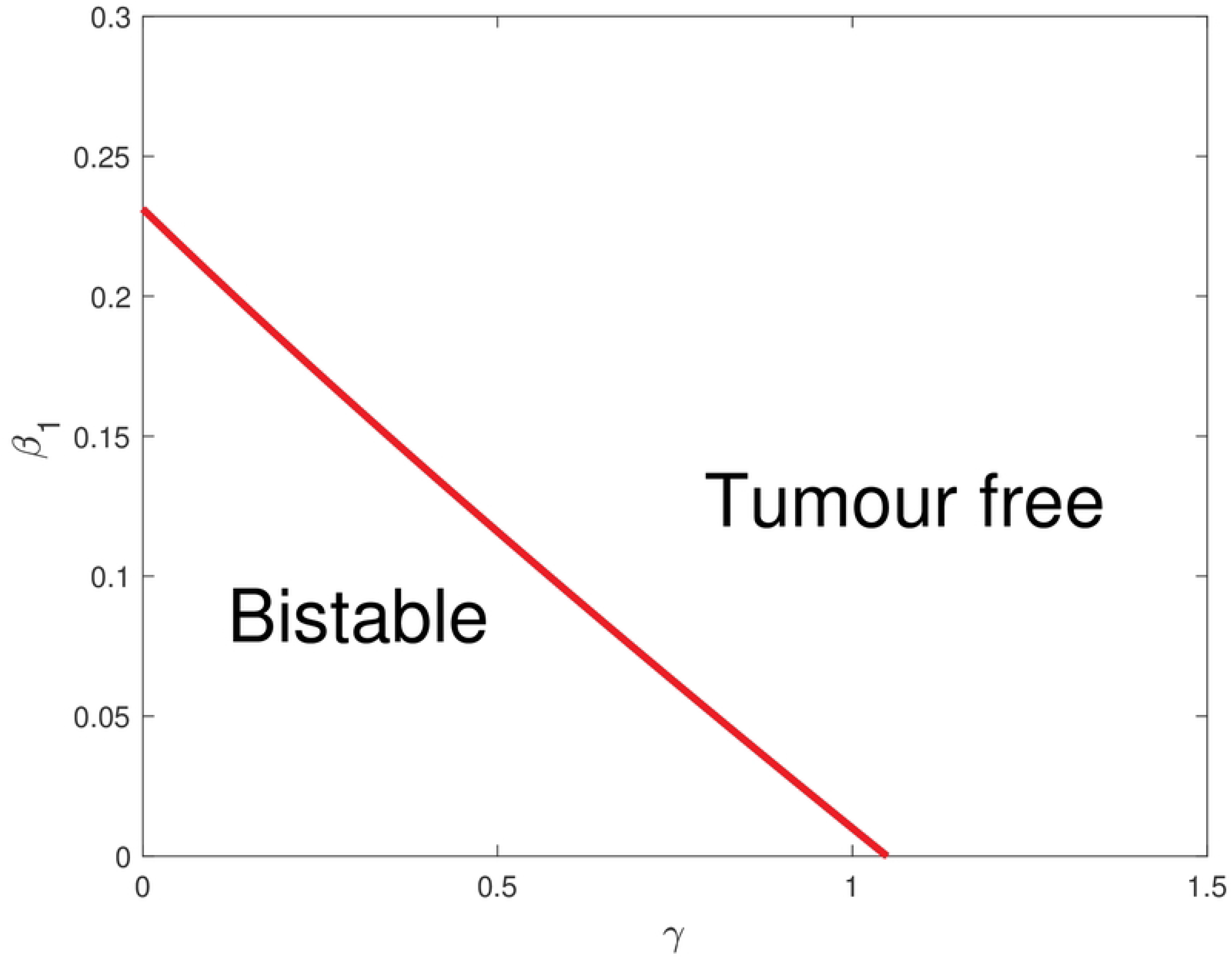

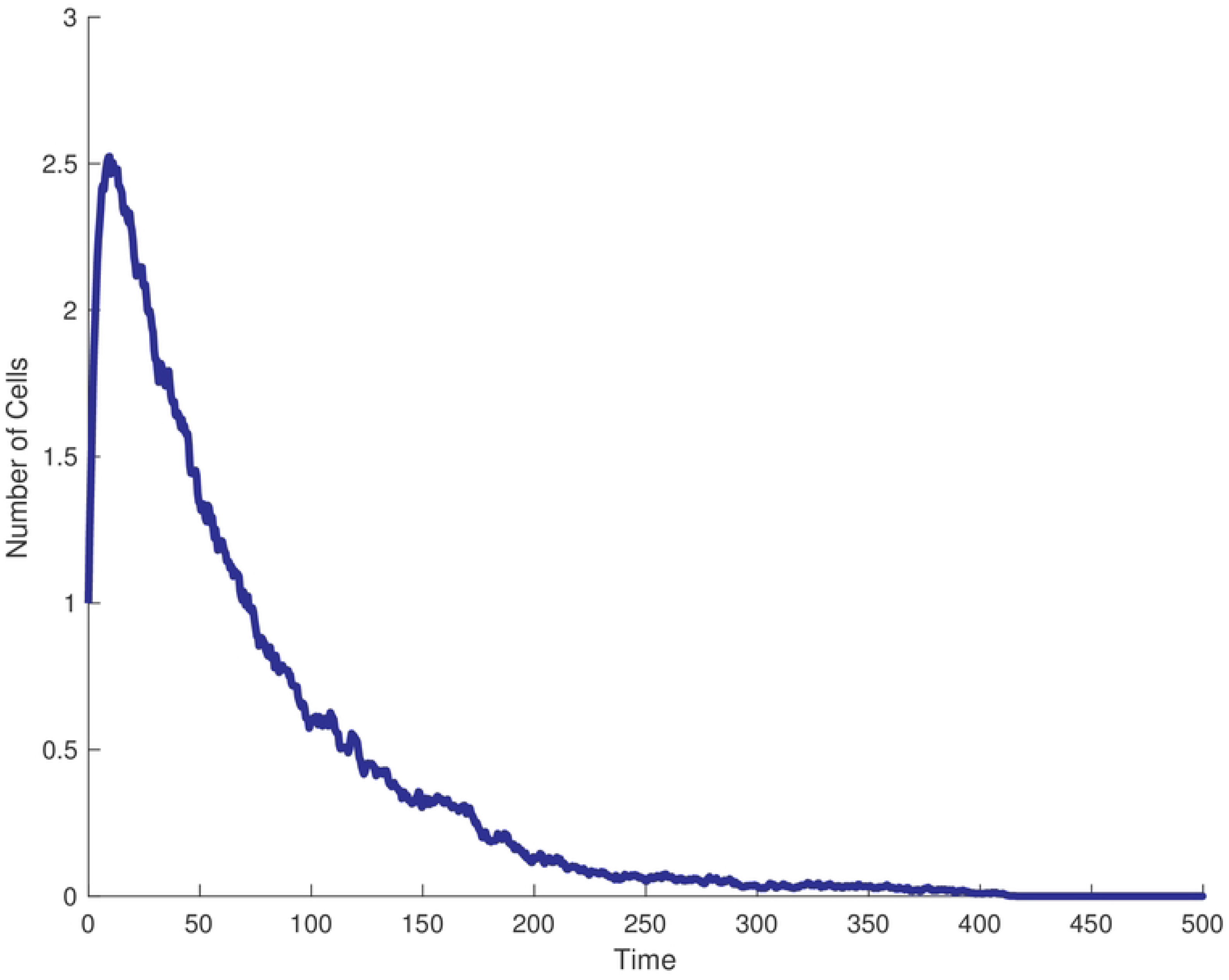

However, when the system is at the bistable region (*γ* = 1 and *β*_1_ = 0), the noise pushes the system into the tumour-free steady state. The system (due to the intrinsic noise) crosses the separatrix of the unstable steady state and settles into the stable lower steady state. This occurs even if we start close to the higher stable steady state branch. Two conclusions can be drawn from this finding. First, the average number of tumour cells increases as time increases, and then reaches a critical value before decreasing to the lower steady state. Second, the system with intrinsic noise always trends to the lower stable steady state. This may indicate that the tumour-free state is the favourite basin of attraction for the system.

#### 2.4 Noise affecting the maximum

As previously discussed, the mean of the number of the tumour cells shows a pulse-like response. This pulse response is usually associated with a negative feedback loop in the dynamical system [34]. Thus, the reason for this pulse-like response can be explained as follows. When the tumour cells undergo rapid proliferation, there is a time delay for the immune cells to be activated in order for them to eliminate the tumour cells. Once the number of immune cells reaches a critical value, they are able to eliminate the tumour cells. This forms a pulse-like behaviour in the dynamic of the tumour cells with a maximum threshold. Therefore, we are interested in how this maximum is affected by some critical parameters, in particular how increasing vitamin intake and chemotherapy, namely *γ* and *β*_1_, affects the maxima threshold. We will focus our discussion on the maximum value and its location. This allows us to estimate the time required for the immune cells to be reach a critical level in order to kill the tumour cells. Fig (7) acts as a reference in our comparison (*γ* = *β*_1_ = 0). First, we shall discuss the effects of increasing *β*_1_ on the pulse-like response. The result is shown in Fig (7). When *β*_1_ = 0, the peak of the average number of tumour cell is ≈ 4 and the peak occurs when t ≈ 18 (see the vertical lines). However, increasing *β*_1_ reduces the peak and reduces the time to reach the maxima of the curve. The biological interpretation of this is that increasing the treatment accelerates the inhibition of the tumour proliferation and hence reduces the number of the affected cells. However, this is not the case when *γ* is increased. As shown in Fig (7), there is no significant changes in the maxima and its location when increasing the vitamin intake.

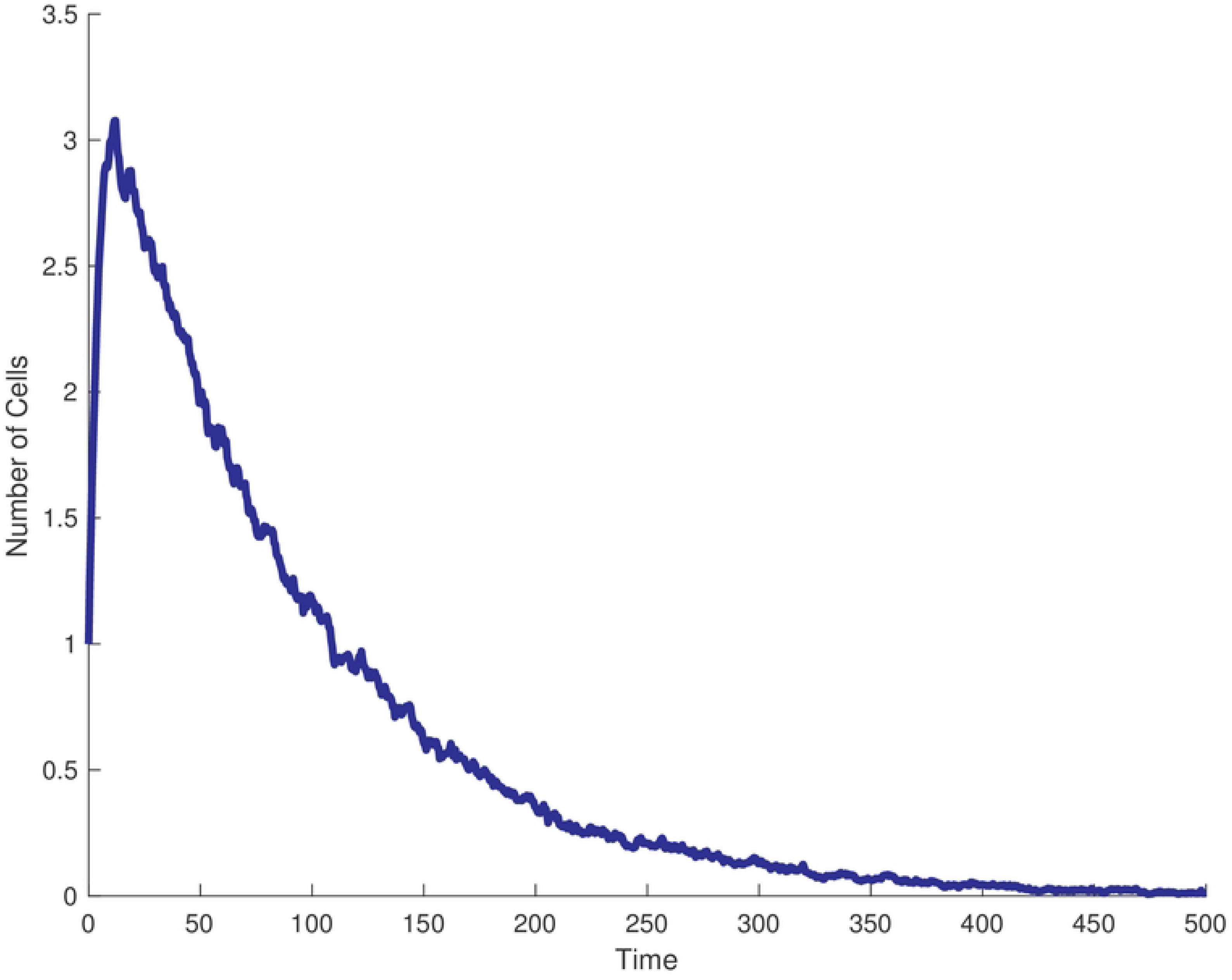

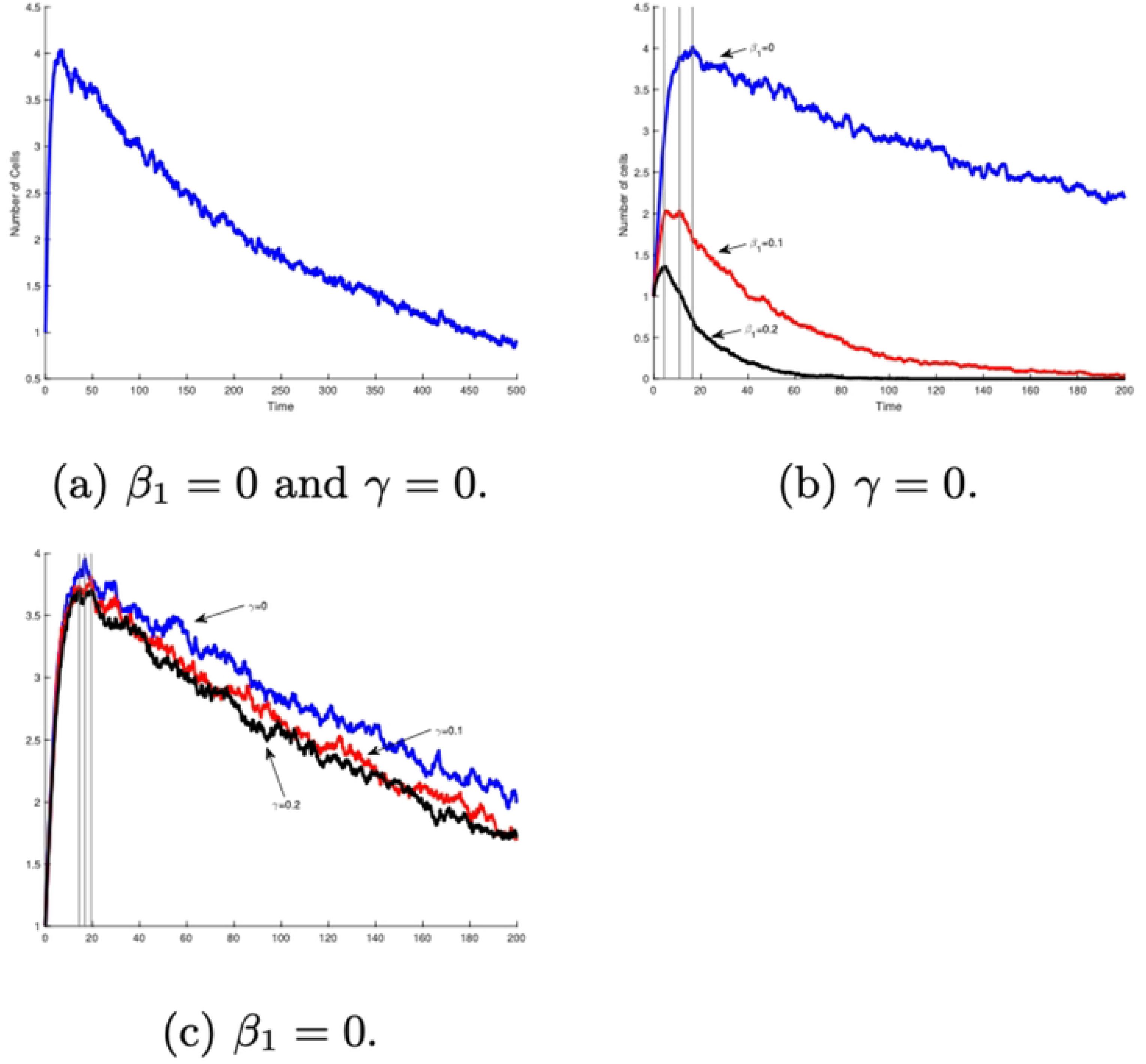

### 3 Mean of elimination time for tumour as a function of *β*_1_ and *γ*

In tumour treatment, it is vital that the cancer cells are driven to extinction. Moreover, the time to extinction is also crucial [11]. In this section, we calculate the average time for a mature tumour to be eliminated from the host. In a previous study [47], they calculated the ratios of various effectors to target cancer cells as a function of time, and showed that as this ratio increases, the elimination time decreases. We followed a different path in our own research. We calculated the mean time of the tumour elimination as a function of chemotherapy dose *β*_1_ and vitamin intake *γ*. This allowed us to investigate whether or not the elimination time can be improved by varying these parameters. In our investigation, we applied Mean First Time Passage (MFTP). We set the system into the active tumour state, and ran 10,000 simulations using the SSA to calculate the average time the system requires to eliminate the tumour. As illustrated by Fig (8), when *β*_1_ i is low, the MFPT is high, and when *β*_1_ is increased the MFPT is reduced. This shows that a sufficient level of treatment can accelerate the transition between higher tumour state to a tumour-free state. However, this is not the case when the vitamin intake is increased. Fig (8) shows that the mean of the elimination of time as a function of the vitamin intake, *γ*. This figure illustrates that the mean is slightly improved as *γ* is increased. However, the effects of increasing *β*_1_ i is stronger.

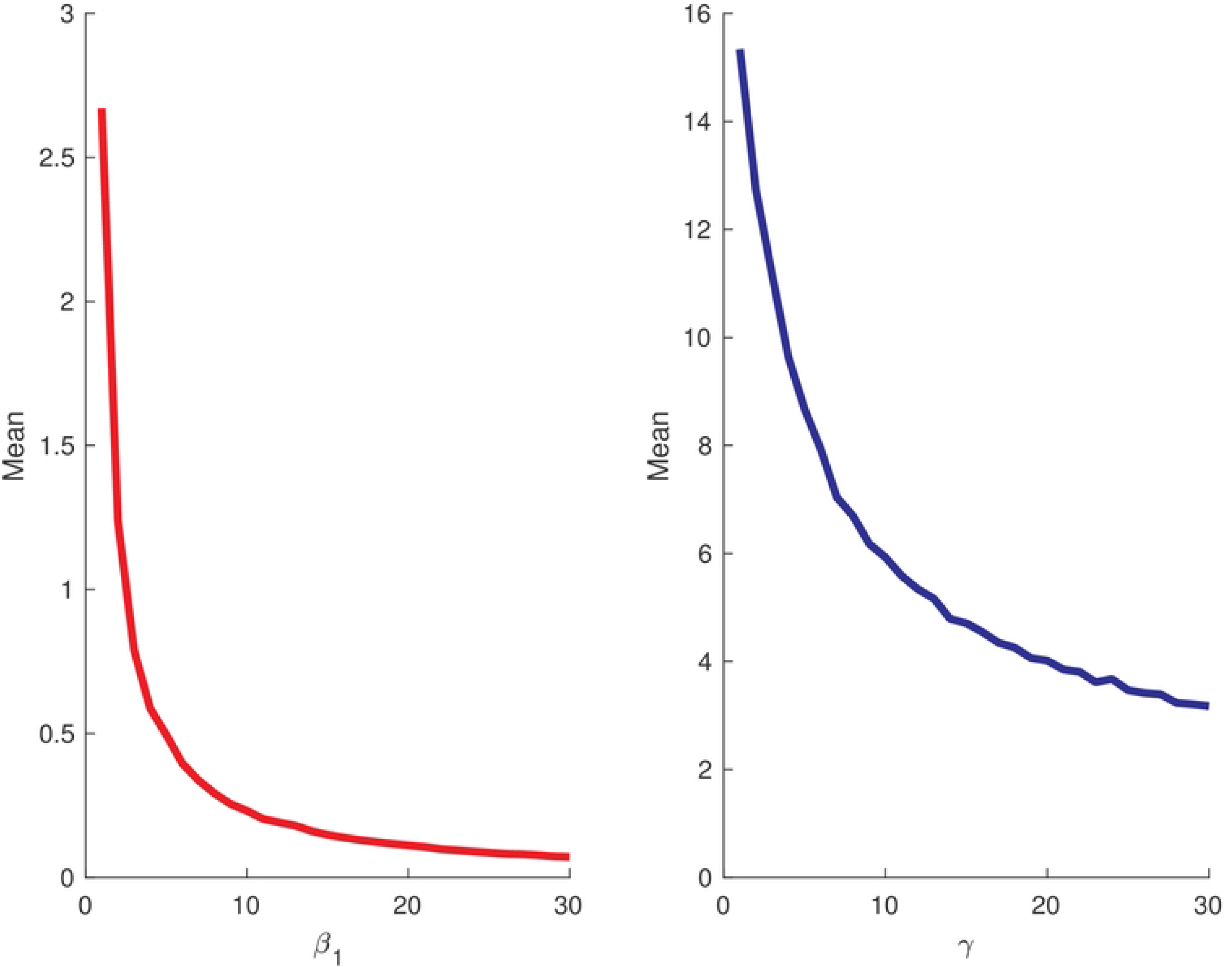

#### 3.1 How does the vitamin affect the dominance durations of the tumour cells

Stochastic simulation allows us to investigate the effect of the dominance duration of the number of tumour cells on the number of the immune cells. We define the dominance duration as the time that the concentration or the number of tumour cells exceeds the number of the immune cells. We shall investigate the duration time of the tumour and immune cells and how they can be altered by increasing *γ*. To calculate the duration time, we ran 10,000 trajectories using the SSA and the result is shown in Fig (9). The blue and red colours represent the tumour and immune cells, respectively. As can be seen from this figure, the blue curve reaches a certain level before declining to its steady state. Moreover, the number of tumour cells stays higher than the number of immune cells. At the time (indicated by the arrow) the number of tumour and immune cells are equal. We investigated how the dominance durations of the tumour cells are altered by the increase in the vitamin intake when *γ* = 0.1, 0.2 and *γ* = 0.5. and shows in panels (a)-(d), respectively. As the vitamin intake is increased, the dominance duration of the tumour cells is significantly decreased. The number of cancer cells reached its maximum in less time than the case when there is no vitamin intake (see Fig (9) (d)). Moreover, the immune cells reach its steady state level faster when there is sufficient vitamin level.

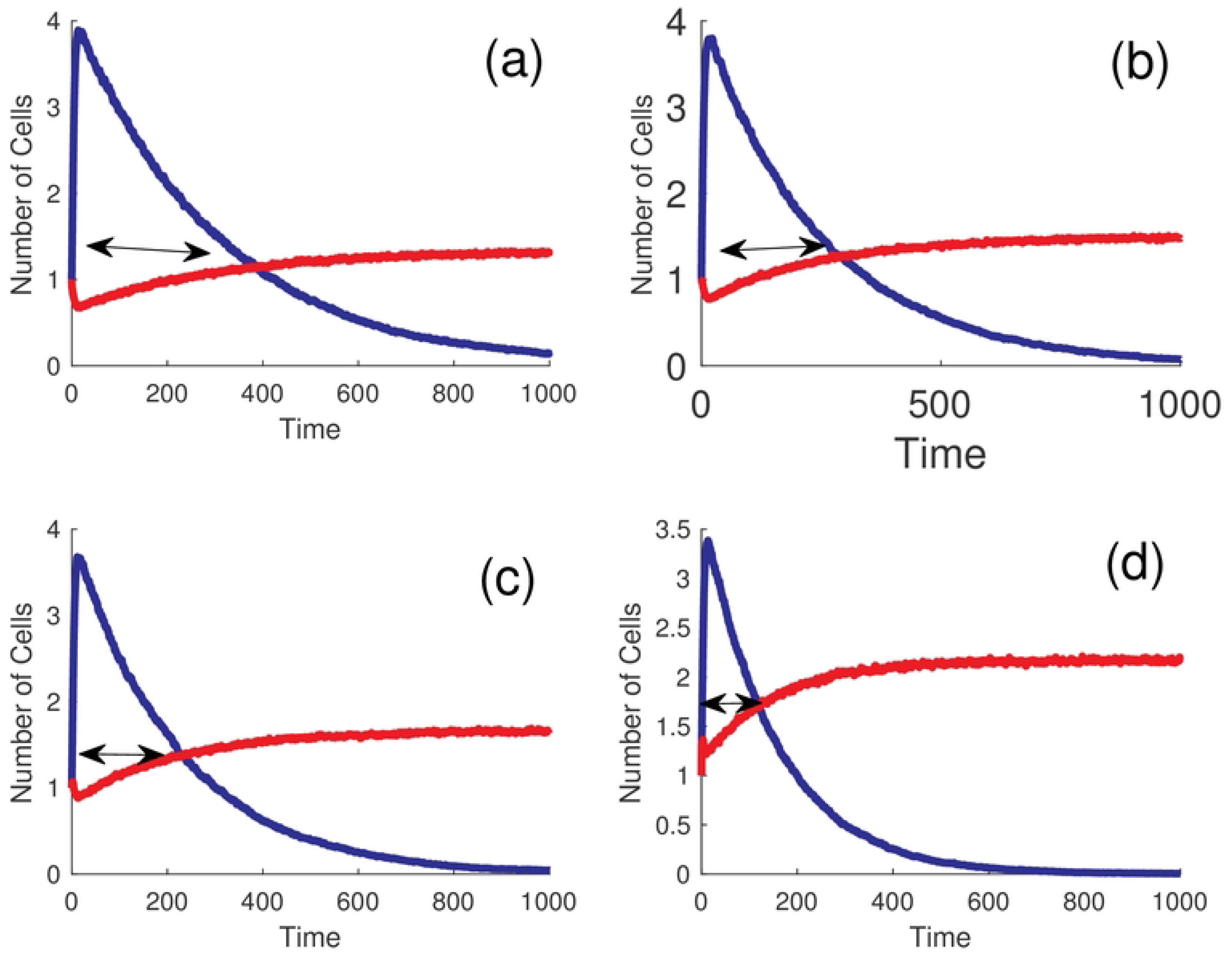

### 4 Deterministic vs stochastic steady state

After discussing the effects of noise on the deactivation time and how can they alter by various parameters, we shall now investigate the impacts of the noise on the steady state level and compare it with its corresponding deterministic model. This allows us to determine whether or not the deterministic model can capture the full behaviour of the system. In particular, we focus on if the deterministic model is able approximate the stochastic average. To do so, we consider four different cases to assess our comparison, namely when *γ* = *β*_1_ = 0 which forms the reference for our discussion. Then, we examine the case when *β*_1_ *>* 0 and *γ* = 0. Thirdly, we discuss the case when *γ >* 0 and *β*_1_ = 0. Finally, we consider the case when *β*_1_ *>* 0 and *γ >* 0.

#### 4.0.1 No chemotherapy and no vitamin intake

When *γ* = *β*_1_ = 0, there is no treatment and no vitamin intake is given to the patient. The result is shown in Fig (10). The blue and red curves show the stochastic trajectories for the cancer and immune cells, respectively. The blue and red dashes lines represent their corresponding deterministic solutions. As illustrated by this figure, the deterministic and stochastic trajectories are slightly different. The deterministic solution could not well approximate the stochastic average.

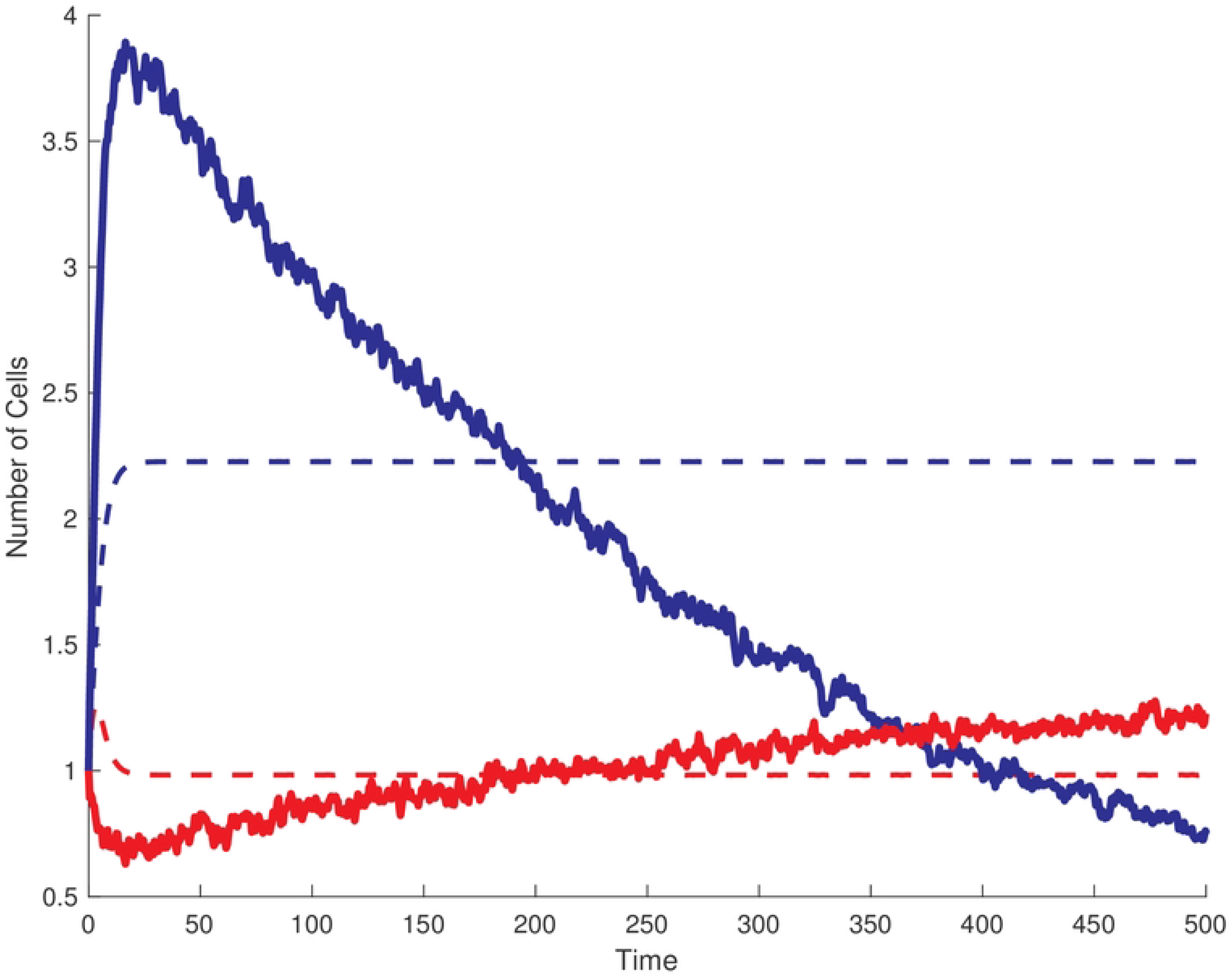

#### 4.0.2 Treatment without vitamin

Next we considered the case when there is some level of treatment without any vitamin intakes. As show in Fig (11), the deterministic model for the cancer cells could not approximate the stochastic average. In fact, the stochastic model drops to lower steady state level than the deterministic model. However, the stochastic solution for the immune cells is well predicted by the simple deterministic model.

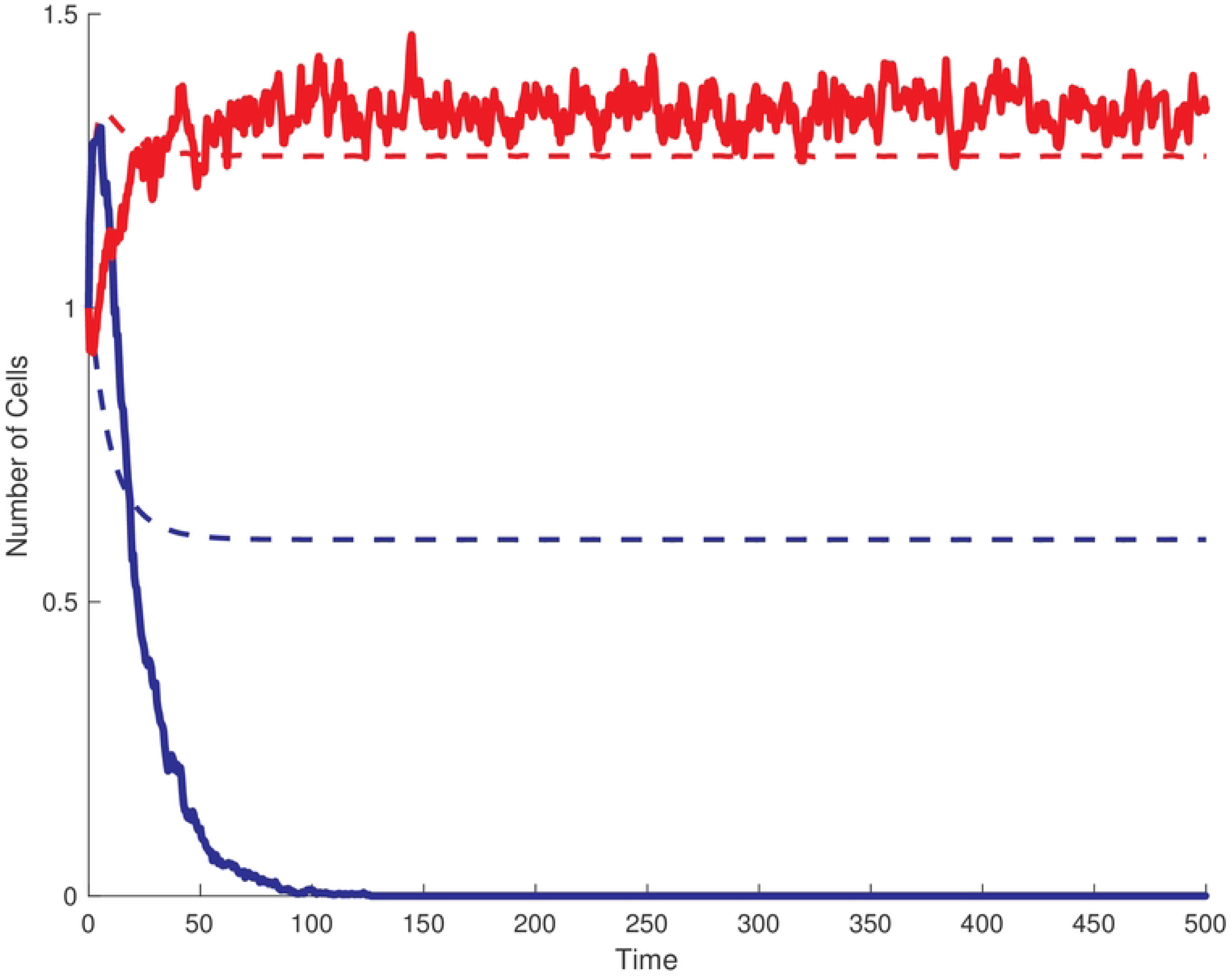

#### 4.0.3 Vitamin without treatment

Fig (12) shows the case when we post the immune cells with some vitamins and there is no treatment is provided to the patients (*γ* = 0.2 and *β*_1_ = 0). As shown in this figure, the deterministic model could not predict the stochastic averages for the cancer and immune cells. In the absence of any treatment plan, the stochastic and deterministic models reach different steady state levels, unlike the previous case when *β*_1_ *>* 0 when the stochastic and deterministic models reach the same steady state for the immune cells.

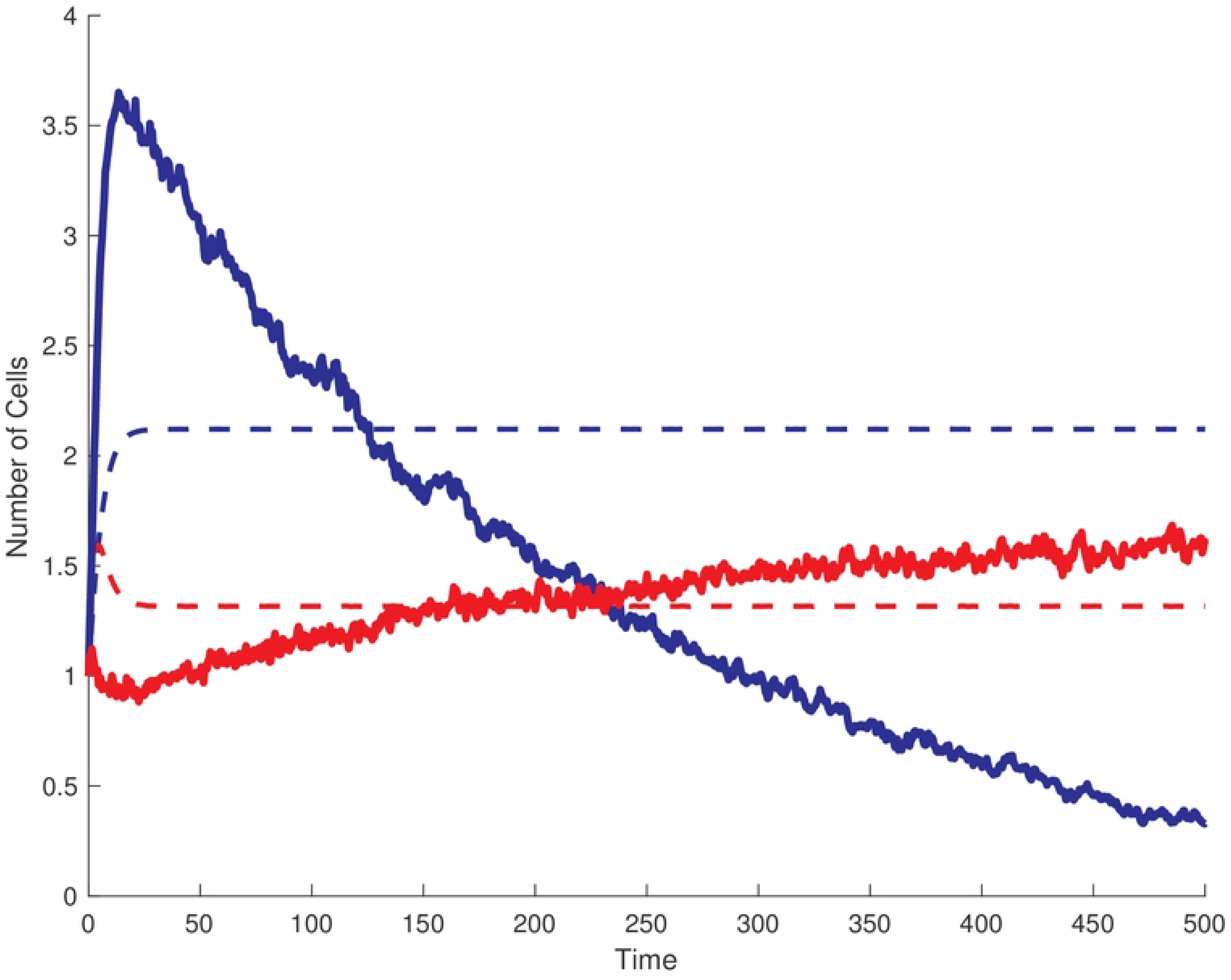

#### 4.0.4 Vitamin and chemotherapy

Finally, we consider the case when there is both treatment and we post the immune cells with vitamins, shown in in Fig (13). The deterministic steady state and stochastic average are identical. The stochastic average in this case is well approximated by the simple deterministic model. The deterministic model predicts the two steady state levels for both the cancer and immune cells.

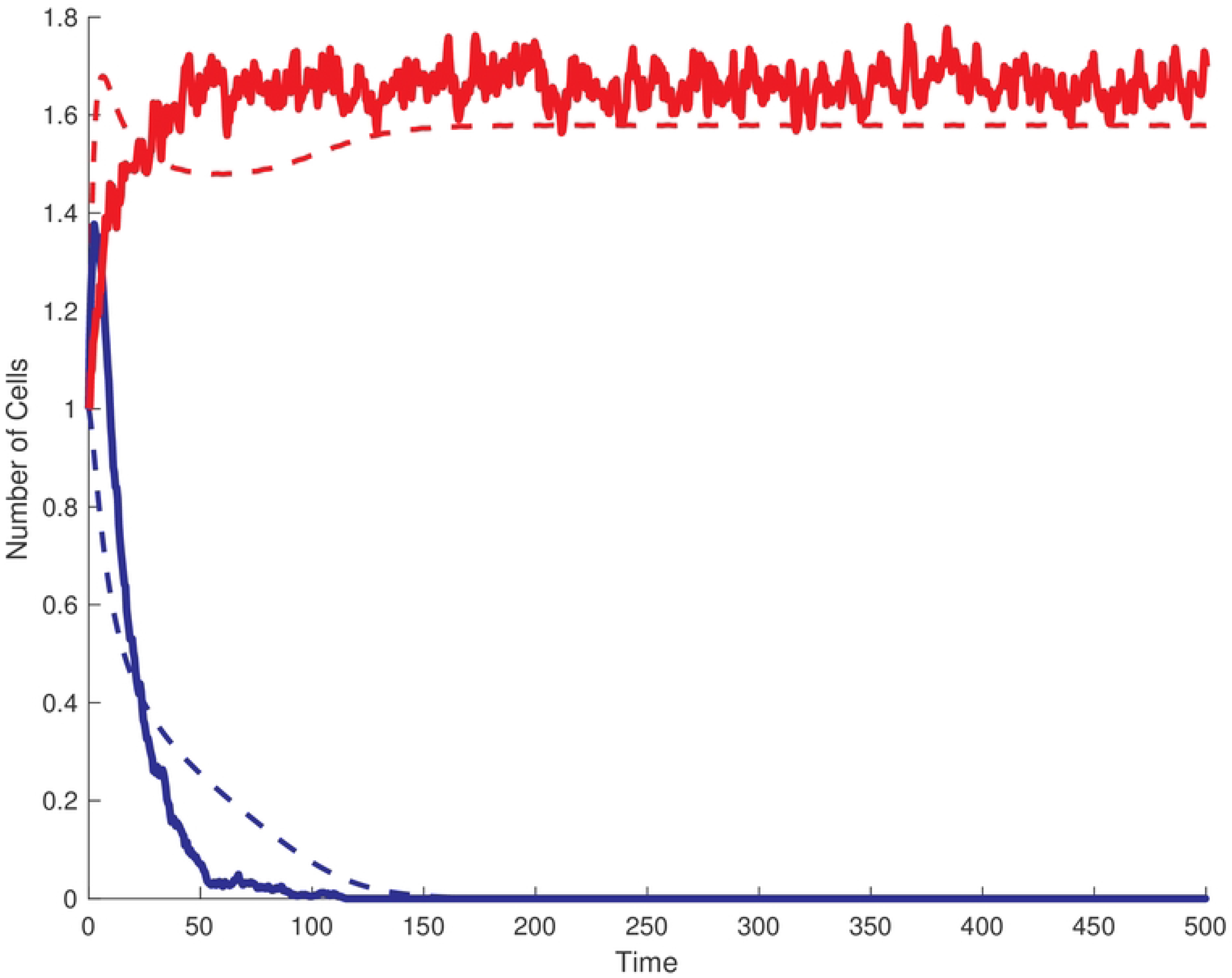

Based on the analysis provided, the following conclusions can be drawn. Firstly, without any treatment, the deterministic model fails to accurately predict the stochastic means. However, in the second scenario where only *β*_1_ *>* 0, both the deterministic and stochastic models reach a consistent steady state level for the immune cells. Lastly, in the fourth case where *β*_1_ *>* 0 and *γ >* 0, both the deterministic and stochastic models converge to the same steady states for both the immune and cancer cells.

#### 4.0.5 Memoryless property

In the bistable region, the system exhibits two stable states which are separated by an unstable one. We monitored the transition from the upper steady state to the lower stable branch using the FTP approach. We ran 10,000 stochastic simulations using SSA. During the simulation, we set the system into the high tumour steady state and the simulation is stopped once the system reached a tumour-free state. Our investigation focused on the transition from the upper steady state to the lower stable branch. The distribution of waiting times for this transition is depicted in Fig (14). The waiting time distribution follows an exponential pattern and exhibits a memoryless property. This suggests that the transition between these two stable basins of attraction is independent of their previous history. In other words, the immune cell does not retain any memory of when it last eliminated a cancer cell.

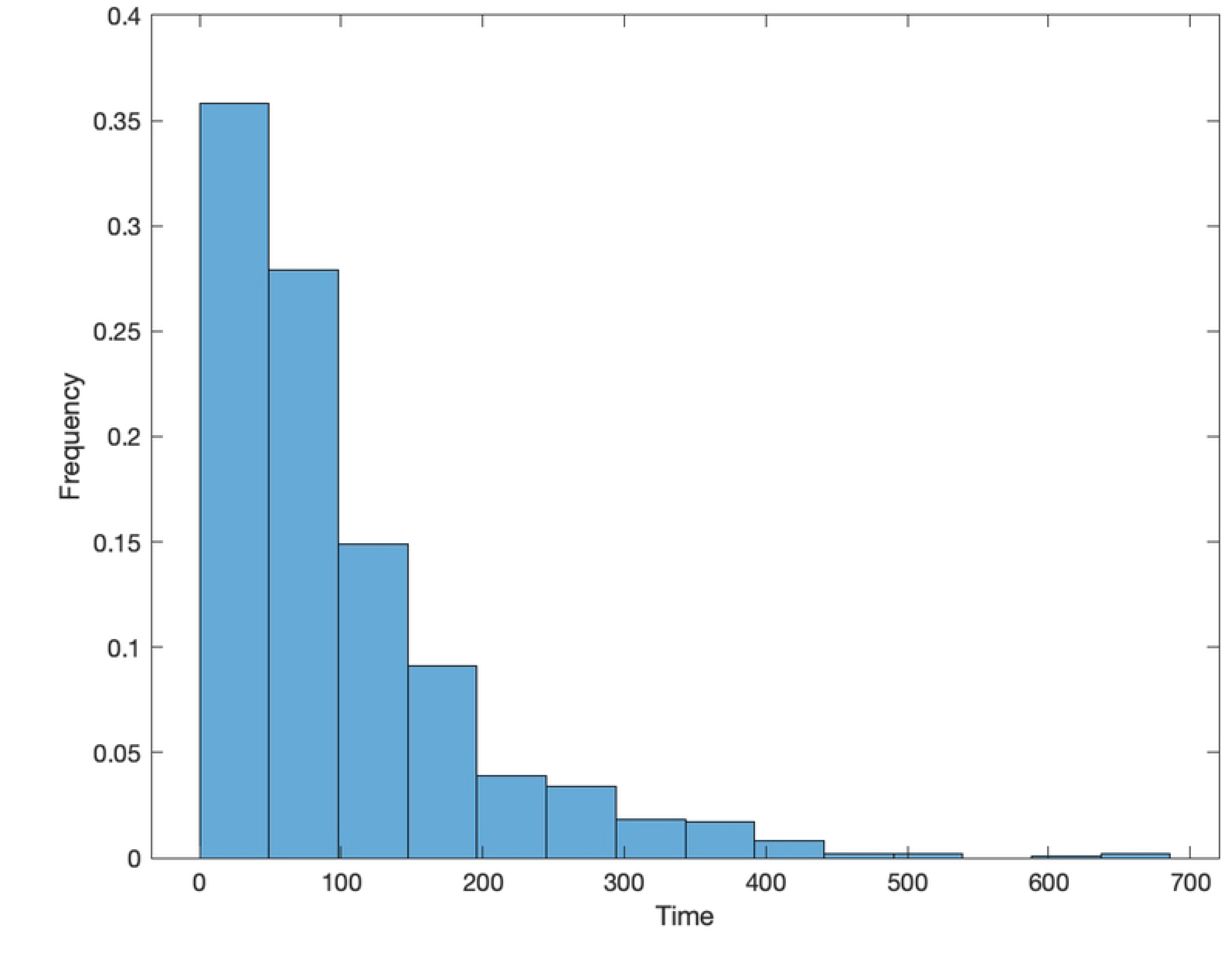

## 5 Conclusion

In this paper, we have discussed the effects of noise on the dynamics of the tumour and immune system interaction. The original paper concluded that the immune system cannot eliminate the tumour cells with proper treatment. This was examined in a high number of the tumour cells using a deterministic approach. However, this is not the case when the population of the cells is low - the immune system can eliminate the tumour at its early stage. The original work calculated the threshold required to eliminate the cancer from the host. Here, we focused on the time required for the system to eliminate the tumour cells. We showed that the waiting time for the elimination of the cancer cells is exponentially distributed. Thus, it indicated memoryless properties where the elimination of any tumour cell does not depend on the waiting time of eliminating its neighbour cell. We also illustrated that the dominance duration time of the tumour cell is reduced when increasing treatment. Finally, the low copy number of the population destroys the bistable switch that was generated when the population of the cells is sufficiently large.

